# NLoed: A Python package for nonlinear optimal experimental design in systems biology

**DOI:** 10.1101/2021.06.03.446189

**Authors:** Nathan Braniff, Taylor Pearce, Zixuan Lu, Michael Astwood, William S. R. Forrest, Cody Receno, Brian Ingalls

**Affiliations:** Department of Applied Mathematics, University of Waterloo, Waterloo, ON, N2L 3G1, Canada

## Abstract

**Motivation:** Modelling in systems and synthetic biology relies on accurate parameter estimates and predictions. Accurate model calibration relies, in turn, on data, and on how well-suited the available data is to a particular modelling task. Optimal experimental design (OED) techniques can be used to identify experiments and data collection procedures that will most efficiently contribute to a given modelling objective. However, implementation of OED is limited by currently available software tools that are not well-suited for the diversity of nonlinear models and non-normal data commonly encountered in biological research. Moreover, existing OED tools do not make use of the state-of-the-art numerical tools, resulting in inefficient computation.

**Results:** Here we present the NLoed software package. NLoed is an open-source Python library providing convenient access to OED methods, with particular emphasis on experimental design for systems biology research. NLoed supports a wide variety of nonlinear, multi-input/output, and dynamic models, and facilitates modelling and design of experiments over a wide variety of data types. To support OED investigations, the NLoed package implements maximum likelihood fitting and diagnostic tools, providing a comprehensive modelling workflow. NLoed offers an accessible, modular, and flexible OED tool-set suited to the wide variety of experimental scenarios encountered in systems biology research. We demonstrate NLOED’s capabilities by applying it to experimental design for characterization of a bacterial optogenetic system.

**Availability:** NLoed is available via pip from the PyPi repository; https://pypi.org/project/nloed/. Source code, documentation and examples can be found on Github at https://github.com/ingallslab/NLoed.

**Supplementary information:** Supplementary materials are available online.

## 1 Introduction

Biological systems are heterogeneous, in terms of both their components and the interactions between them. Mathematical modelling of biological systems provides researchers with a valuable toolset for investigating this complexity. Models can be used to generate new hypotheses about extant systems or to predict properties of novel synthetic biological designs. These models are typically nonlinear, multi-dimensional, and dynamic, and depend on parameters that cannot be directly measured. Accurate estimation of the values of these parameters is critical: the accuracy of the parameterization determines the utility of the model predictions and the model’s overall value as a tool for investigating system behaviour [Hagen et al., 2013]. Accurate parameterization of complex models often faces challenges due to the high cost of data collection. This challenge can be exacerbated by uncertainty in how to design experiments to maximize calibration accuracy.

Recent progress in experimental techniques, specifically with respect to automation and high throughput methods (reviewed by Braniff and Ingalls [2018]), reduce the barriers imposed by experimental costs. However, without complementary advances in experimental design, cheaper data collection may lead to increases in the quantity of data, but not in its quality, and so may have limited impact on model calibration accuracy. Experimental design tools are especially important for nonlinear models, for which intuition can be a poor guide. The value of optimal experimental design in improving parameter estimates for nonlinear and dynamic models in system biology has been previously demonstrated [Hagen et al., 2013, Apgar et al., 2010].

Optimal experimental design (OED) techniques were originally developed in statistics for fitting regression models accurately with minimal experimental effort [Franceschini and Macchietto, 2008]. These techniques have been expanded to nonlinear and dynamic models, and have seen increasing use in the past decades, including applications to biological systems [Kreutz and Timmer, 2009, Chakrabarty et al., 2013, Braniff and Ingalls, 2018]. However, most of these methods rely on custom implementations of their specific numerical algorithms. The lack of established software tools has no doubt limited the use of OED in experimental studies, confining the use of OED techniques to research groups with specialized knowledge of OED methods (e.g. Bandara et al. [2009], Ruess et al. [2015]).

Existing optimal experimental design tools include statistical software suites such as SAS and R, which provide a number of optimal design packages [Groemping, 2020, Atkinson et al., 2007]. These packages are primarily aimed at regression-type models used in statistics and are of limited use to systems biologists.

Several software packages for OED have emerged from the pharmacokinetics-pharmacodynamics (PKPD) research community; examples include the PopED package from Nyberg et al. [2012] (available in MATLAB) and PFIM 3.0 from Bazzoli et al. [2010] (available in R). For a full comparison of OED tools in the PKPD field see Nyberg et al. [2015]. These PKPD tools focus on nonlinear and dynamic models. However, PKPD methods emphasize mixed-effects models (with test-subject-specific sources of variability), which can be computationally demanding and are less relevant for those working outside the PKPD field.

Software development for Bayesian optimal design has been rare, possibly due to the heavy computational cost. A very early example is Clyde [1993]; a more recent example is the `aceBayes` package in R (Overstall and Woods [2017]), which has seen some application to dynamic biological models [Overstall et al., 2020]. Bayesian techniques offer great flexibility in modelling uncertainty and avoid the parameter dependence of local optimal design methods. However, this flexibility comes at additional computational costs resulting from the need for extensive Monte Carlo sampling or other forms of numerical integration.

Wider adoption of OED by experimental systems biologists can be facilitated by software tools that are focused on the specific needs of these researchers. Specifically lacking are optimal design tools — and model building tools in general — for data that is not Gaussian distributed. Systems biology is replete with experimental assays that have non-Gaussian distributions, including plate counts (Poisson), gene expression (log-normal) and viability assays (Bernoulli or binomial). The majority of published OED approaches rely on Gaussian approximations which are often only appropriate under restrictive assumptions.

Also lacking are software tools that enable the iterative experimentation that is needed for modelling nonlinear biological systems — especially in a dynamic context. Nonlinear systems can exhibit dramatic differences in behaviour over parameter and input ranges. *A priori*, experimenters may have a poor understanding of which regime is relevant for their study; iterative experimentation is generally required to identify the operating regime and focus on the dominant effects. While some amount of iteration could ideally be reduced by using Bayesian design tools such as *aceBayes*, there remain logistical reasons why iteration is needed in experimental workflows – the added computational cost of Bayesian designs can also make rapid iteration difficult. NLoed differentiates itself from the existing tools by focusing on providing an easy-to-use and efficient iterative experimental design workflow for nonlinear and non-Gaussian systems.

## 2 Methods

In this work we present the NLoed software package, a purpose-built and user-friendly OED software toolset developed to make OED more accessible to systems biologists. The NLoed package has been released under an open-source licence; it is hosted on Github (https://github.com/ingallslab/NLoed). The package is written in Python 3 and can be used either through a Python interpreter or in Python scripts. The NLoed package is object-oriented; it provides classes for model building and experimental design. The package classes and their associated methods can be flexibly interfaced to facilitate a variety of workflows for both real experiments and simulation studies. NLoed uses Pandas and Numpy data structures, allowing easy data exchange with other numerical packages.

In developing NLoed, we aimed to implement well-established local optimal design methods that are simple and practical, while making use of state-of-the-art numerical methods. The OED methods in NLoed rely heavily on the Fisher information matrix for design optimization. Calculation of the Fisher information relies, in turn, on accurate and rapid local parametric sensitivity computation. Moreover, design optimization requires a nonlinear programming solver. Although it is possible to estimate sensitivity and objective derivatives as finite-differences, these methods can be computationally costly and inaccurate [De Pauw and Vanrolleghem, 2006]. As an alternative, NLoed relies on CasADi [Andersson et al., 2019], a rapid prototyping package for optimal control that provides automatic differentiation functionality and a direct interface to the nonlinear programming solver IPOPT [Wachter and Biegler, 2006]. CasADi allows for rapid and accurate computation of both the Fisher information and design objective derivatives. CasADi also provides a symbolic interface for model construction, facilitating the efficient formulation of complex models.

All models in the package take the following mathematical form:

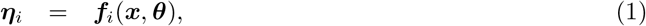

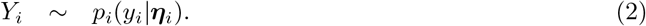

Here the vector ***x*** captures the model inputs while the vector ***θ*** consists of the model parameters. The index *i* is used to distinguish separate observations. For each value of *i*, the function ***f**_i_*(·) maps model inputs and parameter values to the observation’s *sampling statistics*, collected in vector ***η**_i_*. The sampling statistics characterize the distribution that describes a given observation. For example, for a normally distributed observation variable, the sampling statistics are the mean and variance (***η*** = (*μ, σ*^2^) = (mean, variance)), which completely characterize the distribution from which the observation is drawn. (Recall that in statistical modelling, observations are treated as realizations of a random variable that is established by the model; the function ***f***(·) describes that random variables in terms of the model inputs and parameters.) In the NLoed package, the function ***f***(**·**) is implemented using CasADi’s symbolics, which can consist of analytic expressions or numerical algorithms. The number of observations and inputs an NLoed model can accept is constrained only by the computational cost of computing optimal designs. NLoed can be used to optimize designs for multi-input/output models, including multi-state dynamic systems.

Equation 2 specifies each observation, *Y_i_*, as drawn from the probability distribution specified by the sampling statistics ***η**_i_*. The notation *P_i_*(*y_i_|**η**_i_*) emphasizes the conditioning on the sampling statistics. Using this model formulation, NLoed accommodates experimental observations that are non-normally distributed, including counts and strictly positive data. The user can specify a specific distribution type for each observation variable according to the experimental scenario. Supported distributions include Normal, Poisson, Binomial, Bernoulli, Lognormal, Exponential, and Gamma. NLoed is thus applicable to a wide variety of experimental scenarios (e.g. involving both gene expression and plate counts) and modelling frameworks (e.g. deterministic approximations of stochastic models via moment closure [Lakatos et al., 2015] or the linear noise approximation [Elf and Ehrenberg, 2003]). Note that an observation *Y_i_* corresponds to a measurement from a specific observation channel (e.g. concentration, light intensity, optical density, etc.) at a specific, pre-specified time-point. (So, e.g., a measurement repeated at two different time-points corresponds to two different observations *Y_i_*.)

For a given system (1-2), an experimental design in NLoed is defined by a pair of sets 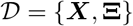 characterizing inputs and observation replicate counts, as follows. Here ***X*** is the set of input vectors, ***x**_j_*, that describe the set of experiments to be executed, i.e. the set of experimental conditions to be observed. The input vector can encode the numerical settings of treatments such as chemical concentrations or light intensity, or environmental conditions such as temperature or media composition. Categorical perturbations, such as strain or treatment type, can be encoded as binary entries. Time-varying inputs are implemented as piece-wise constants by first subdividing the experimental time window and then specifying a constant value over each sub-interval.

The system specification (1-2) defines the collection of possible observations *Y_i_*. The set **Ξ** contains the allocation of replicates: observable *Y_i_* is to be assessed *ξ_i,j_* times under the *j*th input setting. These are (non-negative) integer valued counts. However to improve numerical tractability, NLoed relaxes this integer constraint so that *ξ_i,j_* correspond to non-negative real-valued weights. These real-value allocations are then rounded to discrete values after an optimal solution is found. We refer to a design with real-valued replicate allocations as *relaxed*, while a design with integer-valued allocations is called *exact*.

Design optimization problems in NLoed have the following general form:

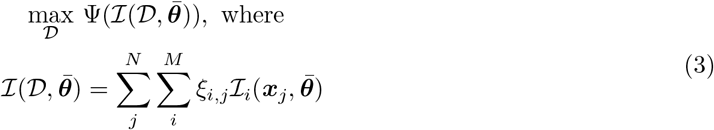

Here 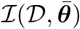 is the expected Fisher information matrix for experimental design 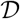, evaluated at the nominal parameter vector 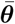. The Fisher information appears in the objective because it is asymptotically related to the expected variability of estimates of the model parameters determined by maximum likelihood estimation [Fedorov and Leonov, 2013]. The Fisher information therefore serves as a useful metric for design utility. The objective function, Ψ(·), maps the Fisher information matrix to a scalar objective. By default NLoed maximizes the determinant of the Fisher information matrix as its objective; this results in a D-optimal design which minimizes the expected confidence volume of the parameter estimates.The integer *N* is the number of unique input vectors considered in the design and *M* is the number of observables (i.e. output channel-timepoint pairs). The relaxed replicate allocations, *ξ_i,j_* are constrained to sum to one: 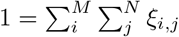

D-optimal design is widely used due to its utility and efficiency, but it cannot be applied to non-identifiable models (for which the Fisher information matrix may be singular, with zero determinant across all design candidates). NLoed does not currently provide objectives for these more challenging scenarios, but it will alert users to singularity issues that could be used to guide model reduction or the adoption of alternative designs.

The expected Fisher information for an overall design is the sum of individual Fisher information matrices for each observation, assuming the observations are statistically independent. (Currently NLoed only supports optimal design for experiments with independent observations.) NLoed then computes the expected Fisher information for an overall design using the sum of individual Fisher information matrices, 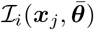, for each input vector, ***x**_j_* and observation, *y_i_*. The individual Fisher information matrices are defined as

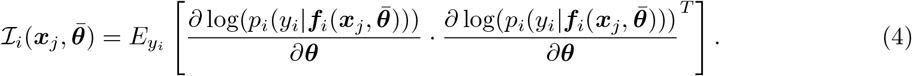

This is the expected value of the square of the parametric sensitivities of the log-likelihood (i.e. variance of the score vector) [Fedorov and Leonov, 2013]. NLoed uses a chain rule decomposition to simplify the computation of the Fisher information – this approach also enables easy computation of the Fisher information for non-Gaussian distributions (see Fedorov and Leonov [2013] for further details.) From the definition in Equation 2 we have that ***η**_i_* = ***f**_i_*(***x, θ***) and therefore the log-likelihood sensitivity vector can be decomposed as:

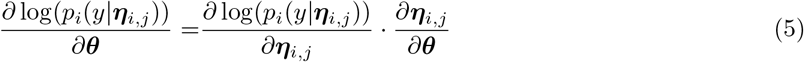

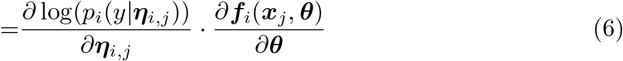

The term *∂**f**_i_*(***x**_j_, **θ***)/*∂**θ*** is the parametric sensitivity of the sampling statistics. This sensitivity can be determined via automatic differentiation of the user-provided model function ***f***(·). The term *∂* log(*p_i_*(*y*|***η**_i,j_*)/*∂**η**_i,j_* and the expectation can be subsumed into the matrix 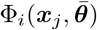, defined as:

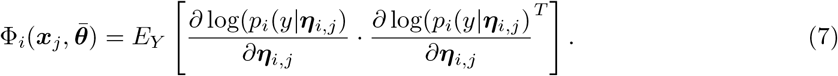

which can be computed analytically from the user-provided function ***f***(·) for the observation distributions supported by NLoed – including several non-Gaussian distributions [Fedorov and Leonov, 2013]. The Fisher information for each input and observation in a design can then be evaluated as

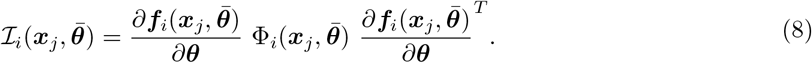

For numerical tractability, the design optimization problem can be further simplified in a variety of ways, depending on how the candidate input vector ***x**_j_* and replicate allocations, *ξ_i,j_* are encoded as optimization variables in the nonlinear programming solver. NLoed offers the users ample flexibility in how the optimization problem is posed via various user-controlled settings.

In addition to design optimization, NLoed includes several model-building and diagnostic tools, including methods for model fitting, sensitivity analysis, model simulation, data sampling and design evaluation. These auxiliary tools are complementary to NLoed’s primary OED functionality. They facilitate incorporation of NLoed into a complete model building and experimental workflow that aligns with the user’s optimal design objectives.

## 3 Implementation

The NLoed library is built around two core classes: the `Model` class, and the `Design` class. The `Model` class captures all of the mathematical information specified in the model framework in Equations (1-2). The `Model` class also provides methods for model calibration and simulation and for evaluating a given design’s performance on the model instance. The `Design` class accepts `Model` instances, as well as other design information, and then implements and solves the design optimization problem. The `Design` class also provides methods for rounding a relaxed design to an exact design with a specified total sample budget. Further details on the class architecture can be found in the documentation provided on the NLoed Github repository.

### 3.1 Model Building

As an example, we consider the CcaS/CcaR optogenetic system described in Schmidl et al. [2014]. (This system was previously characterized in Olson et al. [2017]. A closely related system was characterized in Olson et al. [2014] by a model similar to the one presented below.) We focus on the system’s steady-state response to pulse-width modulated (PWM) green light. (Application to a complementary dynamic model is illustrated in the supplement.) We describe the steady-state response by a Hill function model with normally distributed heteroskedastic observation errors:

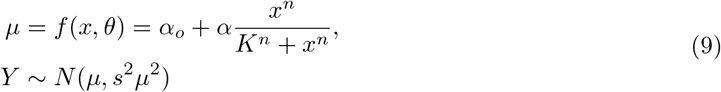

Here the single input *x* is the green light level delivered during growth of the culture (as a percentage of maximal light level). The single independent sampling statistic *η* = *μ* is the observation mean. The single observable *Y*, assumed to be normally distributed, is the steady-state mean GFP expression of a batch culture (see the supplement for details). The components of the unknown model parameter vector *θ* = (*α_o_, α, n, K*) characterize basal expression, maximal induced expression, sensitivity (Hill coefficient), and half-maximal input, respectively. The variance of the GFP observations is assumed to be proportional to the square of the mean, with *s* serving as the proportionality constant. We were not interested in optimizing designs for estimates of *s*; its value was therefore fixed based on initial data (see the supplement for details).

To define a model in NLoed we first specify an expression for the function ***f***(·) mapping inputs to sampling statistics using CasADi’s symbolic types. Code listing 1 shows this process for the model in Equation (9). In Listing 1, a log transformation of the original parameters from Equation (9) is used to ensure the parameter values remain positive during fitting. In the final line of Listing 1, a CasADi function, `func`, is generated to implement ***f***(·) from equation (9). The user can then call the NLoed `Model` class constructor to create the NLoed model instance; this is shown in Listing 2.

**Listing 1:**
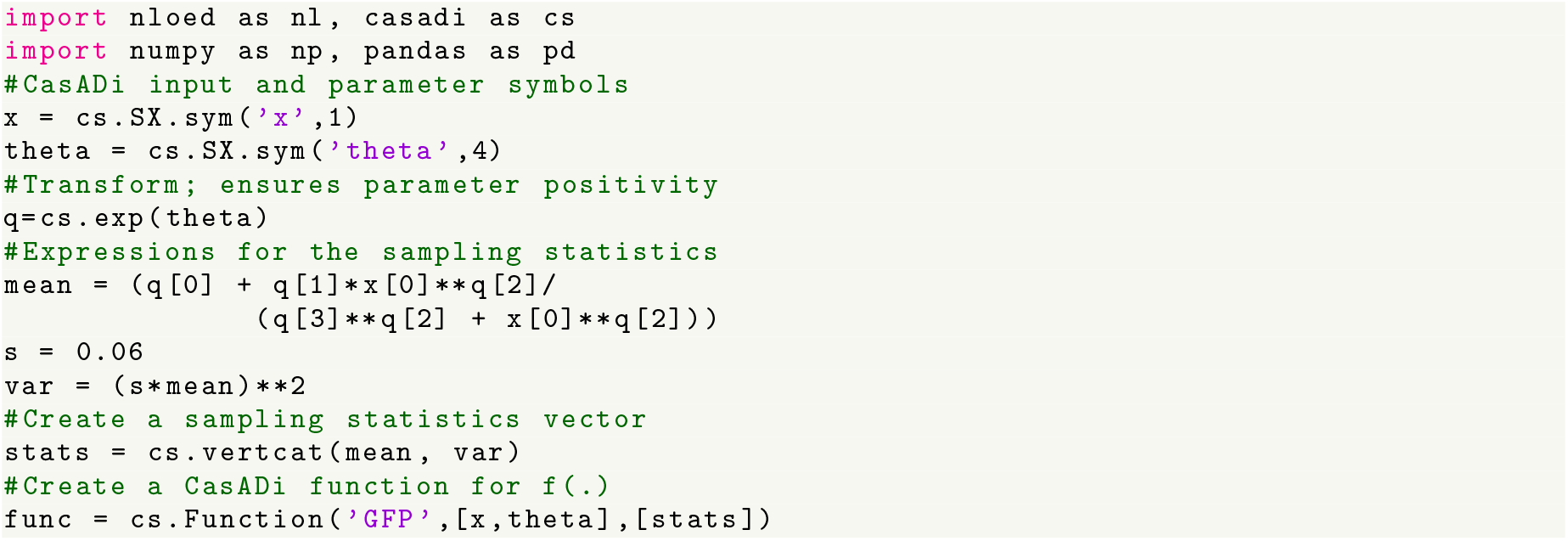
Code for creating a CasADi function for the CcaS/CcaR model’s sampling statistics. The value of *s* set here was estimated from initial data (details in the supplement).

**Listing 2:**
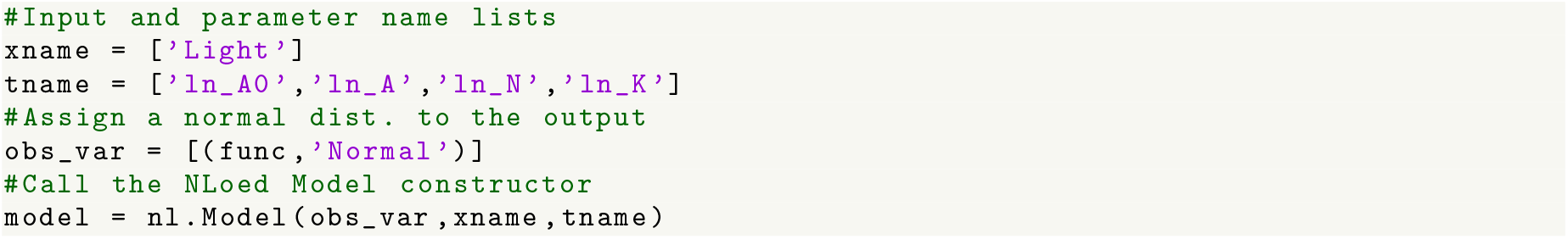
Code for creating an NLoed Model class instance for the CcaS/CcaR model.

For the CcaS/CcaR case study, we conducted initial experiments observing, in triplicate, individual batch cultures’ mean GFP expression in steady-state under 0%, 1.5%, 6%, 25%, and 100% of the maximal green light level (15 observations in total, details in the supplement). This preliminary dataset was used to generate initial parameter estimates, from which NLoed’s locally optimal designs could be determined. Listing 3 details the use of the `Model` class’s `fit()` method in generating initial parameter estimates of *α_o_* = 552.8 a.u., *α* = 9493.2 a.u., *K* = 8.5% and, *n* = 2.4. The variance proportionality constant was estimated independently as s = 0.06 (see supplement for details).

**Listing 3:**
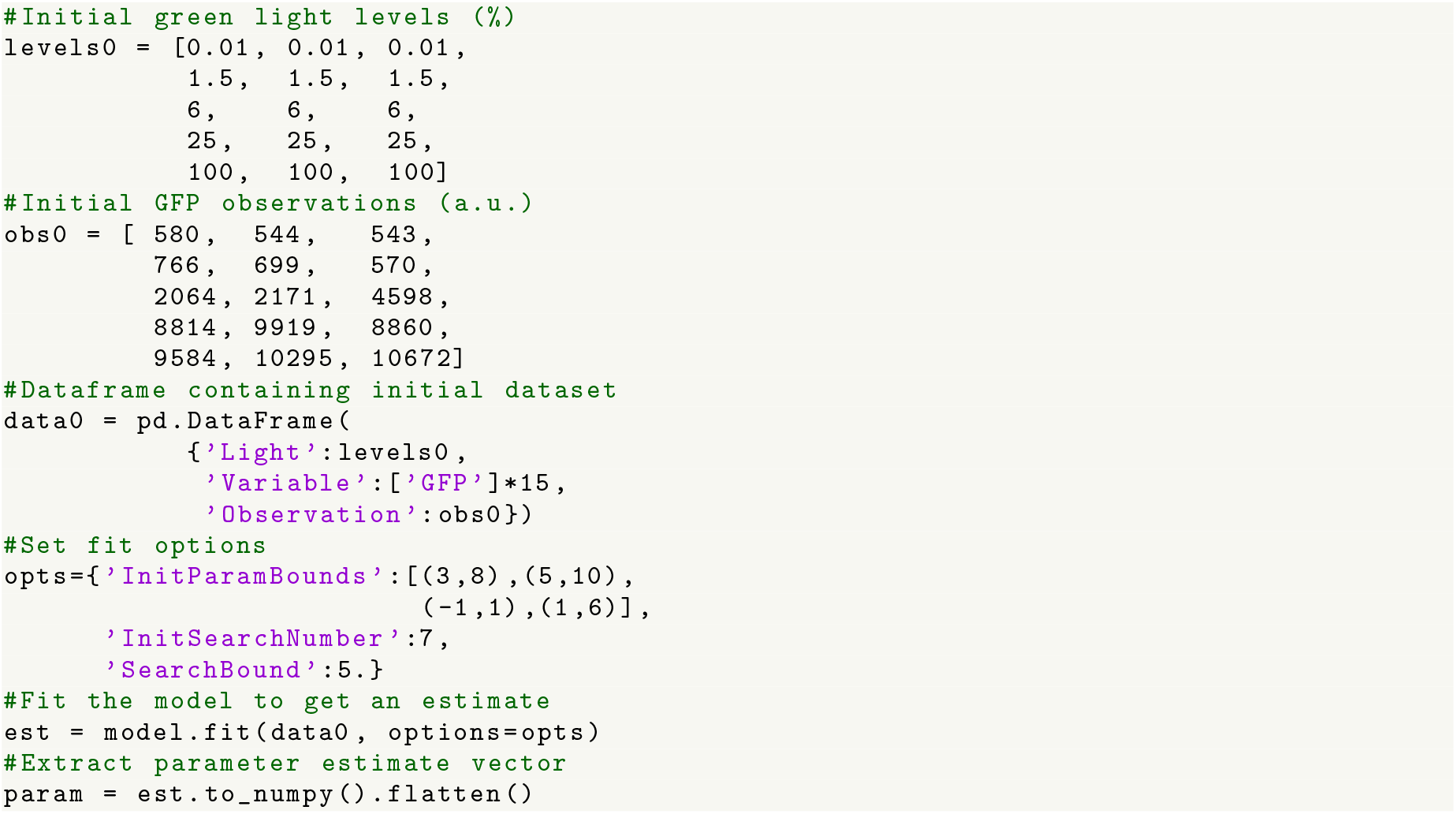
Code for fitting the NLoed Model class instance of the CcaS/CcaR model to an initial dataset.

Beyond fitting, `Model` class instances can perform a number of model development tasks via available class methods:

**Fitting diagnostics:** In addition to implementing a maximum likelihood fitting algorithm (as in Listing 3), the `fit()` method can generate profile likelihood-based confidence intervals for the parameter estimates and can plot visual diagnostics of parameter identifiability such as confidence contours, likelihood profiles and profile trace projections [Bates and Watts, 1988].
**Model predictions:** The `predict()` method allows the user to generate predictions of the means of the model’s observable outputs for a given input setting. This method can also generate prediction uncertainty intervals to quantify the effects of parameter uncertainty on predictions and can perform local parametric sensitivity analysis.
**Design evaluation:** The `evaluate()` method allows the user to evaluate candidate experimental designs with respect to the given model using interpretable metrics such as the expected covariance, bias and mean squared error of the parameter estimates. The `evaluate()` method can use asymptotic or Monte Carlo-based computation to determine the design evaluation metrics.
**Data simulation:** The `sample()` method complements the `predict()` method by simulating random experimental observations from a given experimental design, allowing the user to generate sample data for simulation studies.

### 3.2 Optimal Experimental Design

To generate an optimal design for the CcaS/CcaR model, we passed the previously created `Model` class instance to the `Design` constructor (Listing 4). The call to the `Design` constructor shown in Listing 4 includes specification of the model object, input constraints, design objective, and the nominal parameter values around which the design is to be optimized. In the first lines of Listing 4 the `inp` dictionary is used to define the input constraints. This dictionary specifies the bounds and the number of unique levels of the green light input that are to be allowed during the design optimization. In this case the light level is bounded between 0.01% and 100% and designs consisting of at most four unique levels are considered. NLoed provides a variety of options for how the input levels are handled numerically and which constraints are applied; see the documentation on Github for further details.

**Listing 4:**
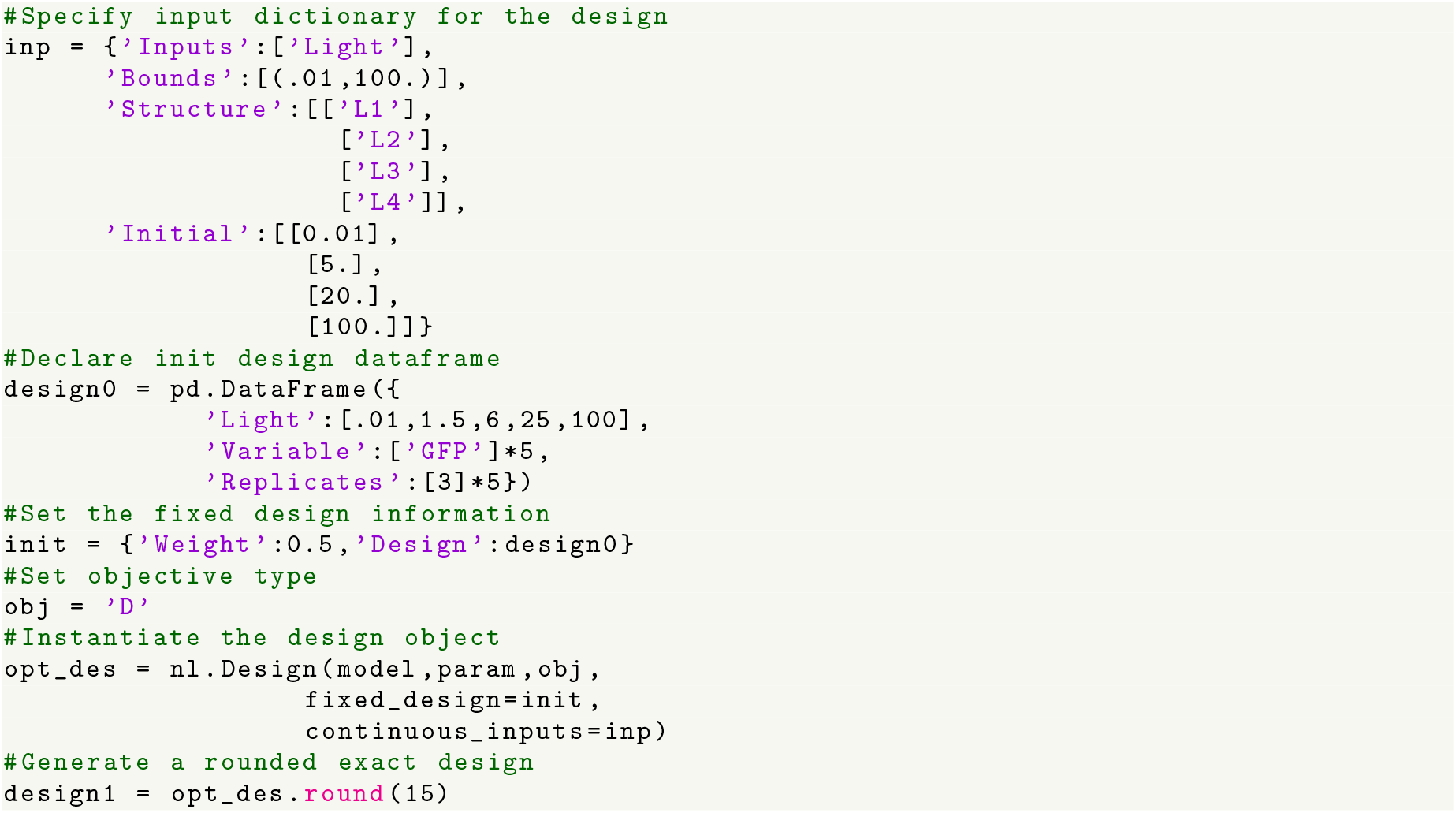
Code demonstrating the creation and optimization of a Design object for the CcaS/CcaR model.

The nominal parameter values are needed in the `Design` constructor call because NLoed uses the expected Fisher information – a local asymptotic approximation – to compute the design objective. The optimality of the resulting design is therefore dependent on how close the nominal values are to the unknown true parameters. It can thus be risky to optimize the design for an entire experiment based on highly uncertain nominal parameter values. This risk can be mitigated by performing a series of sequentially optimized experiments [Atkinson et al., 2007]. In sequential design, rather than using an uncertain nominal parameter set to allocate all observations in a study, the experimenter subdivides their planned experiment into a series of sub-experiments. In this scenario, the experimental design generated for the first sub-experiment may be sub-optimal, but even a sub-optimal experiment yields additional data. This data increases the the sample size – and as parameter estimation accuracy is generally expected to increase with the square root of the sample size [Fedorov and Leonov, 2013] – each additional sub-experiment will on average yield parameter estimates that are nearer to the unknown true value. Therefore, by optimizing each sub-experiment with respect to the current best parameter estimate, the optimality of the combined dataset with respect to the unknown true parameter values is expected to improve. To demonstrate this process, in Listing 4 the parameter estimates from the preliminary experiment are used as the nominal values for the design of an optimized experimental run on the CcaS/CcaR system.

Using the parameter estimates from a previous (sub-)experiment for design optimization will improve the expected performance of the resulting design. However, even greater efficiency can be achieved by conditioning the design optimization directly on past *data* as well. A subsequent experiment can be most efficiently selected if it is optimized to complement the previously-gathered observations with respect to the objective, rather than disregarding the information already available. This approach results in a conditionally optimal design at each iteration of a sequential design procedure: each design is conditionally optimal on all of the past data in the sense that the new design assumes that the past observations will be included in any subsequent parameter estimation. This conditional optimization is shown in Listing 4 where the initial design, `designO`, is passed into the `Design` constructor via the `init` field. By passing in the initial design, NLoed will select a new design that best complements the initial design in order to optimize the objective.

When the `Design` class is instantiated near the end of Listing 4, NLoed runs an IPOPT call to generate an optimal relaxed design, in which the optimal allocation of replicates assumes a continuum of replicate allocation weights Ξ. To produce a design that can be implemented, the last line of listing 4 uses the `round()` method to generate a usable design with a total of 15 replicates (over all observations, mirroring the design of the preliminary experiment). The resulting optimized exact design contained in variable `design1` is shown in Table 1. Note that the optimal design, perhaps non-intuitively, focuses on the lowest part of the input range. This optimal design took NLoed 0.25 seconds to generate on a laptop computer (details in supplement). As a point of comparison, performing a similar optimization with `aceBayes`, a Bayesian OED tool, took 19.179 seconds – about 80 times longer (see supplement for details). We conclude that the local approaches implemented in NLoed can be significantly faster than Bayesian approaches, making them suitable for use in cases when computational costs are limiting (e.g. extensive or rapid iteration, or for analysis of complex models).

**Table 1:** An optimal design for the CcaS/CcaR model.

| Green Light Input | Number of Replicates |
| --- | --- |
| $\mathbf{x}_j$ | $\xi_{i,j}$ |
| $\mathbf{x}_1 = 0.01\%$ | 4 |
| $\mathbf{x}_2 = 2.8\%$ | 5 |
| $\mathbf{x}_3 = 8.1\%$ | 4 |
| $\mathbf{x}_4 = 100.0\%$ | 2 |

We then executed the optimal design and re-fit the model to the combined data (preliminary and follow-up optimal). The updated parameter estimates are *α_o_* = 525.6 a.u., *α* = 9876.1 a.u., *K* = 10.4% and, *n* = 2.3. To compute the size of each parameter’s approximate 95% confidence interval, we used the `Model` class’s `evaluate()` method (Listing 5, full code in the supplement). In Listing 5, the `evaluate()` method produces a parameter covariance matrix for a combined design - including both the initial design, `design0`, and the optimal design, `design1`. The covariance matrix is then used to compute the asymptotic confidence intervals. For comparison, we generated asymptotic confidence intervals for two other cases: (i) the preliminary experiment alone, and (ii) the preliminary experiment followed by a replicate of the preliminary experiment (code provided in supplement). (The replicate initial design provides a controlled comparison for the increased sample size). Figure 1 shows the asymptotic confidence intervals for all three cases. These results show improved precision of the parameter estimates using the optimized design rather than simply replicating the original design. These improvements are primarily accrued in the two nonlinear parameter *K* and *n*, where the optimal design noticeably outperforms direct replication.

**Listing 5:**
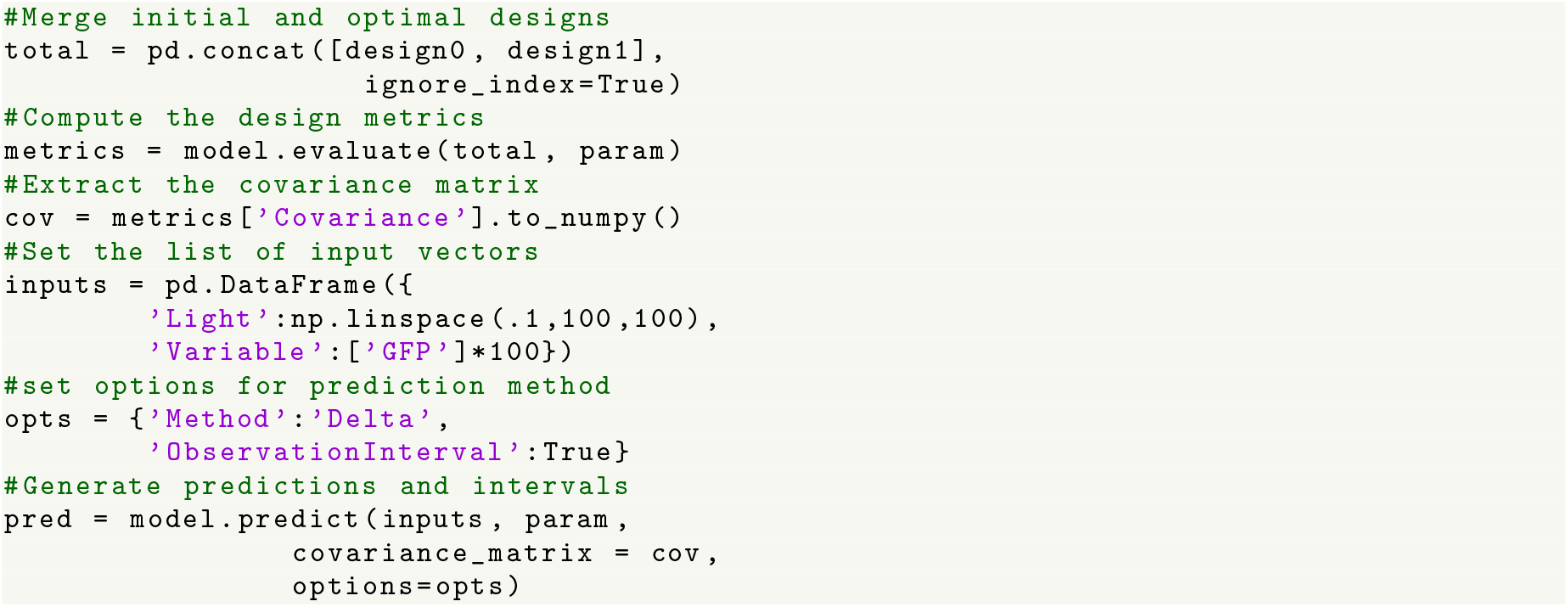
Code for generating design evaluation metrics and model predictions.

**Figure 1:**
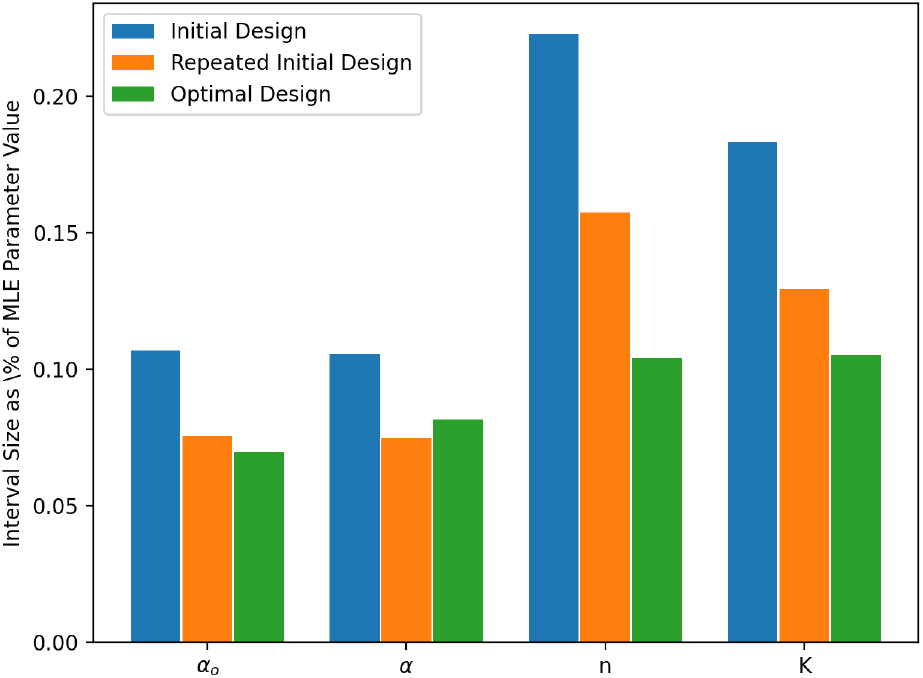
A comparison of 95% confidence interval sizes, expressed as a percentage of the parameter estimate values, between various combinations of the initial and optimal designs.

The impact of the experimental design on model performance can be visualized by the effects of uncertainty on model predictions. NLoed produces the necessary data through the `Model` class’s `predict()` method. After generating the covariance matrix in Listing 5, the matrix is then passed to the `predict()` method to generate model predictions and a 95% uncertainty interval for the observations. This interval accounts for observation variability and the parameter uncertainty resulting from the overall design. Figure 2 shows a plot of the model predictions and observation interval along with the initial and optimal data.

**Figure 2:**
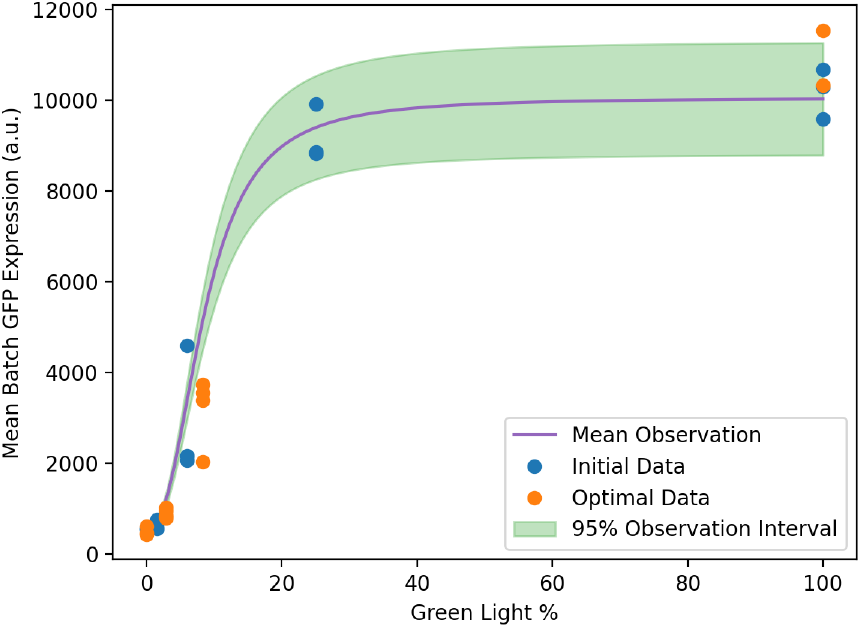
Model predictions and 95% observation interval for the CcaS/CcaR model. Also shown are the initial and optimal data used for fitting the model.

## 4 Discussion

Optimal experimental design (OED) consists of a well established set of methodologies with a long history of success. However, it remains a challenge to translate OED methods into practice, especially for evolving experimental disciplines such as systems biology. NLoed aims to make OED more accessible to systems biologists by providing OED methods in an easy-to-use and open-source package. By combining OED methods with general model building and diagnostic algorithms, NLoed provides a complete modelling workflow. NLoed’s use of state-of-the-art automatic differentiation tools, via CasADi, improves performance and makes the package easily extensible. In summary, NLoed is an ideal tool for rapid iteration of experimental design and model building, where more computationally intensive Bayesian OED methods may not be practical.

Currently, NLoed only implements local asymptotic optimization criteria for parameter accuracy. NLoed is therefore best suited to designing large, iterative characterization experiments. In these cases, when the sample size is large, the asymptotic and local approximations are expected to perform well, especially when used for sequential design. NLoed’s focus on parameter accuracy also means it is especially useful for precise modelling of well understood natural system or synthetically engineered systems where the model structure is reasonably well determined: it is better suited to precision characterization of well studied systems as opposed to early investigation of novel ones.

In future releases we hope to expand NLoed’s capabilities, including new design methods such as pseudo-Bayesian techniques for addressing model and parameter uncertainty [Schenkendorf et al., 2009]. This will improve NLoed’s ability to accommodate uncertainty, especially for early experimental work on novel systems. Regardless, in its current form, NLoed can make it easier for experimentalists to more efficiently allocate laboratory resources while also providing theoretical groups with tools to study the effects of experimental design on model identifiability.

## Acknowledgements

The plasmids used in this work were generously provided by Jeffery Tabor’s group at Rice University, from the work Schmidl et al. [2014].

