## Supplement for "NLoed: A Python package for nonlinear optimal experimental design in systems biology"

### 1 Experimental Materials and Methods

#### 1.1 Strains and Plasmids

All work was carried out in *E. coli* strain BW29655, sourced from the *E. coli* Genetic Stock Center (CGSC) at Yale University. Experimental strains carried plasmids pSR58.6, expressing CcaR and GFP with $Cm_R$, and pSR43.6r, expressing CcaS and other proteins necessary for optogenetic sensing, with $Sm_R$. These plasmids were acquired through Addgene where they were made available by the Tabor group [3]. Both plasmids were transformed into the experimental strain via electroporation. Successful transformation was confirmed via diagnostic digest. The growth media used in this work was M9 minimal media with 20% glucose, 10% casamino acids and antibiotics (50 µg/mL spectinomycin and 25 µg/mL chloramphenicol).

#### 1.2 Growth and Light Exposure

The dose-response protocol for both the initial and optimal experimental designs was as follows:

1. Frozen stock of the BW29655 strain containing both pSR43.6r and pSR58.6 were inoculated into a culture tube containing 1 mL media.

2. The inoculated tube was then grown overnight at 37°C with shaking at 250 rpm on the light array at full red light intensity, zero green light intensity (see below for light array description). Growth was confirmed by spectrometry ($OD_{600}$).

3. The overnight culture was then diluted 1000-fold and returned to the light array to restart growth under full red light intensity and zero green light intensity for 2 hours. Growth was again confirmed by spectrometry ($OD_{600}$).

4. Sixteen experimental culture tubes were prepared by adding 1 mL of the previously described M9 media into each.

5. Each of the sixteen experimental tubes was then inoculated with 100µL of the re-started overnight culture.

6. The experimental tubes were then transferred to the light array and were grown at 37°C with shaking at 250 rpm. Each tube was grown under the prescribed green light intensities for the given experimental design. Red light intensity was maintained at its maximum setting for all experimental tubes.

7. The $OD_{600}$ of the experimental cultures was measured periodically. Cultures were diluted as needed to ensure the $OD_{600}$ remained between 0.01 and 0.2.

8. After 6 hours of growth, the experimental tubes were harvested. From each experimental tube, 100µL of culture was transferred to a microcentrifuge tube and placed on ice.

9. GFP expression was then assayed for a sample from each culture by flow cytometer.

#### 1.3 Flow Cytometry

Analysis was carried out on an Amnis Imagestream MkII flow cytometer (EMD Millipore). Data acquisition was implemented with the Amnis Inspire software package (EMD millipore). All samples were gated against gradient root mean square in the brightfield channel, accepting images with scores between 30 and 80. Fluorescence was excited by a 488 nm solid state laser at 5 mW intensity. Green fluorescence was detected by a 533/55 nm band pass filter. For each experiment, the mean intensity of 10,000 cells is reported.

#### 1.4 Light Array

The light array constructed for this work was inspired by [2]. Each of the sixteen slots in the light array accommodates a single culture tube. Each slot accommodates three LEDs whose intensity can be independently controlled. In the current experiment, red (650nm-Kingbright:WP1503SRD) and green (520nm-Kingbright:WP7083ZGD-G) LEDs were used. The light array was attached to a shaker table inside an air incubator to maintain aeration and temperature control during the experiments. Light intensity on the array is controlled via pulse-width modulation (PWM). The light controller allows the PWM setting to be varied between 0 (no light emitted) and 4095 (maximum output). Images of the light array are shown in Figure 1.

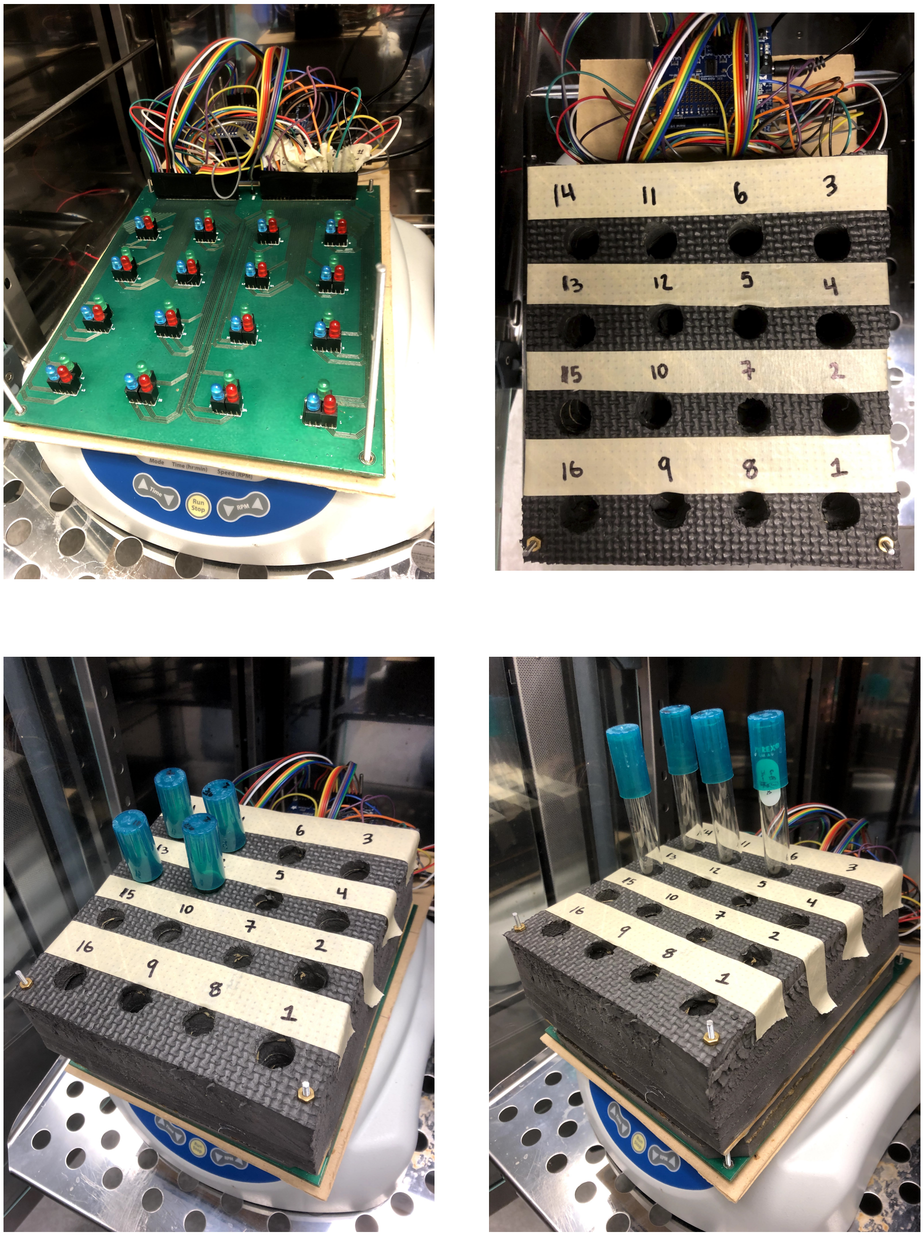


Figure 1: Images of the light array apparatus used in this work. Clockwise from top left; (1) the light array's printed circuit board with LEDs visible – insulating foam for securing test tubes has been removed, (2) view of the unloaded array from above with insulating foam attached and the array locations numbered, (3) view of the array showing its placement on the shaker table with test tubes partially inserted into the insulating foam, (4) a view of the array with test tubes fully inserted.

#### 2 Output Variance

The preliminary experiment involved triplicate measurements in each of five green light level conditions, from which we determined the sample variability for the GFP intensity measurements in each condition (Figure 2). These results revealed significant heteroskedasticity. NLoed can accommodate heteroskedastic error distributions in its maximum likelihood fitting and optimal design in a number of ways, including having the variance depend on variance-specific parameters or having it be proportional to the mean. For this simple case study, we assumed a proportionality. We established a line of best fit (with a zero intercept) to the data in Figure 2. The slope is 0.06, which is the proportionality constant used for design and fitting in the main text.

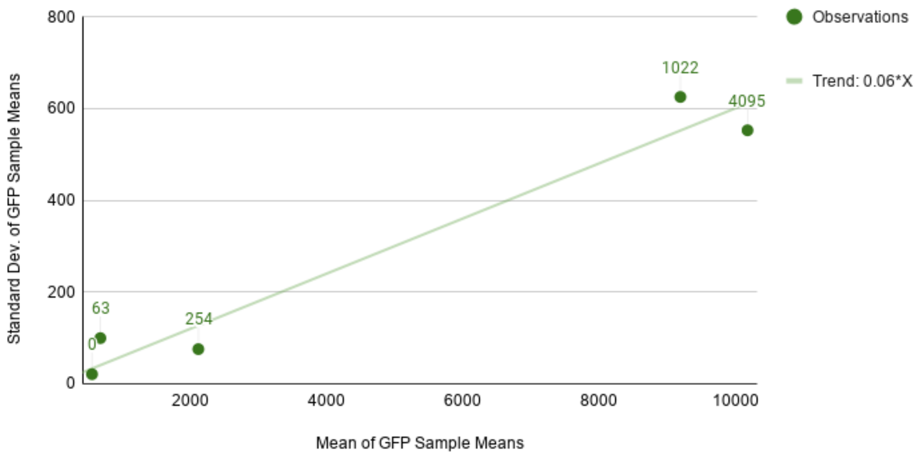


Figure 2: Standard deviation of GFP measurements plotted against the mean of the GFP measurements in each light condition for preliminary experiment. A trend line has been fit to provide a rough approximation for the model heteroskedasticity. One of the triplicate means taken at the 254PWM condition was removed as an outlier.

#### 3 Code for Main Text Figures

The Python code segments used to generate Figure 1 (confidence interval comparison), and Figure 2 (model prediction intervals) from the main text are provided below.

```python
import matplotlib.pyplot as plt
#optimal green light levels
levels1 = \
[.001]*4+[2.88]*5+[8.33]*4+[100]*2
#optimal GFP observations (a.u.)
obs1 = [433,  477,  441,  604,   1032,
        823,  849,  792,  954,   3555,
        2039, 3384, 3740, 10321, 11534]
#dataframe containing initial dataset
data1 = pd.DataFrame(
            {'Light':levels0+levels1,
            'Variable':['GFP']*30,
            'Observation':obs0+obs1})
#fit the model
est = model.fit(data1, start_param=param)
#extract parameter estimate vector
param = est['Estimate'].to_numpy().flatten()
```

Listing 1: Code to refit the model to the combined initial and optimal datasets. The resulting parameter estimates are used to compute the confidence intervals and model predictions shown in the main text.

```python
import matplotlib.pyplot as plt
import scipy.stats as st
#compute  # of std. deviations for a 95% interval
thresh = st.norm.ppf(0.95)
#compute the covariance for the initial design
covariance_init = model.evaluate(design0, param)
#compute the standard parameter error for initial design
stderr_init = np.sqrt(np.diag(covariance_init['Covariance']))
#compute upper/lower confidence bounds for initial design
upper_bnd_init = param + thresh*stderr_init
lower_bnd_init = param - thresh*stderr_init
#combine the initial and optimal designs
opt_design_tot = pd.concat([design0, design1], ignore_index=True)
#compute the covariance for the combined design
covariance_opt = model.evaluate(opt_design_tot, param)
#compute the parameter std. error for the combined design
stderr_opt = np.sqrt(np.diag(covariance_opt['Covariance']))
#compute the upper/lower confidence bounds for combined design
upper_bnd_opt = param + thresh*stderr_opt
lower_bnd_opt = param - thresh*stderr_opt
#replicate the initial design for the duplicated scenario
rep_init_design_tot = pd.concat([design0, design0], ignore_index=True)
#compute the covariance for the replicated initial design
covariance_rept = model.evaluate(rep_init_design_tot, param)
#compute the parameter std. error for replicated initial design
stderr_rept = np.sqrt(np.diag(covariance_rept['Covariance']))
#compute the upper/lower confidence bounds for replicated design
upper_bnd_rept = param + thresh*stderr_rept
lower_bnd_rept = param - thresh*stderr_rept
```

Listing 2: Code for determining the asymptotic confidence intervals of the initial, repeated initial, and the initial and optimal combined designs.

```
#exponentiate the final parameter estimates to convert
#from log scale
exp_vals = np.exp(param)
#exponentiate the confidence bounds for each scenario and compute
#the normalized size of the intervals in the original scale
exp_diff_opt = (np.exp(upper_bnd_opt)-np.exp(lower_bnd_opt))/exp_vals
exp_diff_init = (np.exp(upper_bnd_init)-np.exp(lower_bnd_init))/exp_vals
exp_diff_rept = (np.exp(upper_bnd_rept)-np.exp(lower_bnd_rept))/exp_vals
#set the width of the bars
width = 0.25
#create plot x labels names
labels = [r'$\alpha_o$',r'$\alpha$',r'n', r'K']
#set the x axis settings
x = np.arange(len(labels))
# create the figure
fig, ax = plt.subplots()
#plot the initial design CI interval sizes
rects1 = ax.bar(x - width-0.01, exp_diff_init.flatten(), width, label='Initial Design')
#plot the duplicated initial design CI interval sizes
rects2 = ax.bar(x, exp_diff_rept.flatten(), width, label='Repeated Initial Design')
#plot the combined (optimal + initial) design CI interval sizes
rects3 = ax.bar(x + width+0.01, exp_diff_opt.flatten(), width, label='Optimal Design')
#set y axis title
ax.set_ylabel('Interval Size as \% of MLE Parameter Value')
ax.set_xticks(x)
ax.set_xticklabels(labels)
ax.legend(loc=2)
fig.tight_layout()
#generate plot
plt.show()
```

Listing 3: Code for generating Figure 1 of the main text showing the confidence intervals from the initial, optimal, and combined data.

```python
#create a plot
fig, ax = plt.subplots()
#plot observation interval
ax.fill_between(pred['Inputs','Light'],
                pred['Observation','Lower'],
                pred['Observation','Upper'],
                alpha=0.3,
                color='C2',
                label='95% Observation Interval')
#plot mean model prediction
ax.plot(pred['Inputs','Light'], pred['Prediction','Mean'], '-',color='C4',label='Mean Observation')
#plot initial dataset
ax.plot(levels0, obs0, 'o', color='C0',label='Initial Data')
ax.plot(levels1, obs1, 'o', color='C1',label='Optimal Data')
ax.set_xlabel('Green Light %')
ax.set_ylabel('Mean Batch GFP Expression (a.u.)')
ax.legend(loc=4)
plt.show()
```

Listing 4: Code for generating Figure 2 of the main text showing the data, model predictions and the prediction uncertainty.

### 4 Implementation of Dynamic Models in NLoed

NLoed can be used to analyse dynamic models, but this is a somewhat more lengthy process compared to the algebraic model presented in the main text. Non-linear dynamic models require some form of numerical integration to evaluate model behaviour. The simplest method for implementing such dynamic models in NLoed is to encode the numerical integration procedure itself as a symbolic structure in CasADi symbols. This has the advantage that the entire algorithm can be differentiated via automatic differentiation for FIM computation, objective derivatives and sensitivity analysis. This resulting process, demonstrated below, is rather complex. Moreover, NLoed's computational efficiency when applied to dynamic models depends on the user's choice of numerical integration scheme. Thus greater expertise and optimization may be needed to achieve optimal performance for a given system.

A two-state differential equation model of gene expression is shown below:

$$\begin{aligned} \frac{dx}{dt} &= \frac{\alpha}{1+\frac{K}{u}} - \delta x, \\ \frac{dy}{dt} &= \beta x - \gamma y \end{aligned} \tag{1}$$

Here $x$ and $y$ represent abundance of mRNA and protein product, respectively. Expression is controlled by an inducer with abundance $u$, which acts as an experimental input. The model parameters $\alpha$, $\beta$, $K$, $\delta$ and $\gamma$ are to be estimated. We suppose that the user is planning to perform time series experiments to fit the model and is looking to design such experiments – i.e. to select an inducer profile and measurement schedules for observations of $x$ and $y$ – for accurate parameter estimation.

```
1  import casadi as cs
2  import nloed as nl
3  #create state variable vector
4  states = cs.SX.sym('states',2)
5  #create control input symbol
6  inducer = cs.SX.sym('inducer')
7  #create parameter symbol vector
8  parameters = cs.SX.sym('parameters',5)
9  #log-transformed parameters
10 alpha = cs.exp(parameters[0])
11 K = cs.exp(parameters[1])
12 delta = cs.exp(parameters[2])
13 beta = cs.exp(parameters[3])
14 gamma = cs.exp(parameters[4])
15 #create symbolic RHS
16 rhs = cs.vertcat(alpha*inducer/(K + inducer) - delta*states[0],
17                  beta*states[0] - gamma*states[1])
18 #create casadi RHS function
19 rhs_func =
   ↪  cs.Function('rhs_func',[states,inducer,parameters],[rhs])
```

Listing 5: Code for the creation of a CasADi function for the RHS of the two-state ODE model for gene expression system.

The first step in creating an NLoed model for this gene expression system is to create a CasADi function for the differential equation's right-hand side (RHS). Listing 5 demonstrates this processes. In line 4, symbols are created for the mRNA and protein concentration state variables. In line 6 a symbol is created for the inducer concentration. Line 8 shows the creation of a symbol vector for the model parameters. In lines 10-15, we perform a log transformation (as in the main text) to ensure non-negativity of any fit parameter values. In line 17-18, a symbol for the RHS of the system is created, containing symbolic expressions for Equation (1). In line 20, a CasADi function is created to map the state, inducer, and parameters to the derivatives expressed by the RHS.

```
20 #time step size
21 dt = 1
22 # Create symbolics for RK4 integration
23 k1 = rhs_func(states, inducer, parameters)
24 k2 = rhs_func(states + dt/2.0*k1, inducer, parameters)
25 k3 = rhs_func(states + dt/2.0*k2, inducer, parameters)
26 k4 = rhs_func(states + dt*k3, inducer, parameters)
27 state_step = states + dt/6.0 * (k1 + 2*k2 + 2*k3 + k4)
28 # Create a function to perform one step of the RK integration
29 step_func = cs.Function('step_func',[states, inducer,
   ↪  parameters],[state_step])
```

Listing 6: Code for the creation of a symbolic implementation of a fourth-order Runge-Kutta integrator using the RHS function for the mRNA-protein system.

The CasADi function for the RHS can now be used to create new symbols for numerical time-stepping of the state vector by a given integration method. In Listing 6, imitating examples given in the CasADi documentation, a fourth-order Runge-Kutta algorithm (RK4) is implemented [1]. In line 22, the time step interval is set to 1, and in lines 24-26, the four incremental slopes are computed as CasADi symbols using the RHS CasADi function [1]. In line 28, the incremental slopes are combined to create a symbol for a full time step over the given interval. Finally, in line 30, a CasADi function is created mapping the current state, inducer and parameter values to the next state values, a single time step later.

```
30 #create a symbol for the initial inducer level
31 initial_inducer = cs.SX.sym('init_inducer')
32 #define the steady state initial states in terms of the initial
   ↪  inducer
33 init_mrna = (alpha/delta)*initial_inducer/(K+initial_inducer)
34 ini_prot = beta*init_mrna/gamma
35 # zip the initial states into a vector
36 initial_states = cs.vertcat(init_mrna, ini_prot)
```

Listing 7: Code for symbolic encoding of the initial conditions for the mRNA-protein model's steady states.

The RK4 time-stepping CasADi function, `step_func()`, allows simulation of the system state one step at a time. Because the RK4 stepper is implemented as a CasADi function, we can compute a symbolic vector for the system state at any of these points. This allows considerable flexibility in how a dynamic model is encoded for use in NLOED via the `Model` class. Choices such as whether the initial conditions are handled as known constants, parameters, or as functions of the input can be made based on the experimental situation. The user must also make choices about which state variables will be observable, and the degree of flexibility allowed for input adjustments, sample scheduling and time-series length.

To illustrate, we assign the gene expression system to start at equilibrium with respect to a user-selected inducer level. In Listing 7, lines 32-37, the initial conditions for a given time series run are implemented symbolically. In line 32, a symbol, `initial_inducer`, is created for the initial inducer level. In lines 34-35, the steady-state values of the two state variables are expressed in terms of the initial inducer concentration. In line 37, the initial state variables are concatenated into a single vector, `initial_states`. (In contrast, if the initial conditions were known constants, the `initial_states` vector could be replaced by numerical constants, whereas if the initial conditions were to be fit as parameters, they could be assigned as free CasADi symbols and appended to the parameter vector.)

```python
#3 samples per cntrl interval, 4 cntrl intervals
#control intervals are 1+2+3=6 steps long:
#|ctrl_int1||ctrl_int2||ctrl_int3||ctrl_int4|
#|-1--2---3||-1--2---3||-1--2---3||-1--2---3|
#set number of control intervals
num_cntrl_intervals = 4
#create a vector for inducer levels in each control interval
inducer_vector = cs.SX.sym('inducer_vec',4)
#define a sample pattern to apply in each control interval
sample_pattern= [1,2,3]
#lists to store symbols for each sample point, and times of each sample
sample_list, times = [], []
# set the initial states and initialize the step counter
current_state, step_counter = initial_states, 0
#loop over control invervals
for interval in range(num_cntrl_intervals):
  # loop over sample pattern
  for num_stps in sample_pattern:
    #iterate steps indicated by sample pattern
    for k in range(num_stps):
      #propagate the state variables via integration
      current_state = step_func(current_state, inducer_vector[interval], parameters)
      step_counter+=1
    #save the state symbols and times of each sample
    sample_list.append(current_state)
    times.append(step_counter*dt)
```

Listing 8: Code for symbolic integration to implement a specific sampling and observation pattern and collection of observation variables for various states and time points.

Having defined symbols for the initial conditions, we can now generate symbols for the simulation time-steps using the RK4 stepper function. We will call the RK4 stepper at the initial condition symbols to generate a new symbol for the system state one time step later. The RK4 stepper can then be iteratively called to generate the state symbols at later time steps. The RK4 stepper acts on CasADi symbols for both the inducer level and the parameters and generates symbols for the current state that depend symbolically on the initial conditions, the parameters and the past inducer levels. The user can then record the state symbols at any time points they want considered in the OED analysis.

Each iteration of the RK4 generates a candidate observation time-point. To implement time-varying inputs, the user can assign a value of the control symbol over each time-step, or can lump sequences of time-steps together to produce a coarser set of sub-intervals over which the input schedule is defined. This is illustrated in Listing 8, in which a time series experiment is assumed to have four piece-wise constant control intervals, each of which is subdivided into three candidate observation time-points, at which both state variables can be observed. To propagate the state symbol and record it at at the candidate time points, a nested loop over the control intervals and sample schedule is implemented. In line 43, we set the number of control intervals to be four and in line 45 we create a CasADi symbol vector, `inducer_vector`, containing symbols for the four inducer levels in the time series. In line 47 an array specifies the number of time steps to advance before taking an observation within each control interval. This results in candidate observation points at one, two and three time steps after the beginning of each control interval. In line 49, we create lists to store CasADi symbols for the observed state at each candidate time point as well as the times at which these occur. (Recording the times is useful for naming observation variables in the NLoed model because each time point is its own observation variable in the NLoed `Model` instance.) In line 51, we initialize the system state to the steady state values in the `initial_state` variable and set the step counter to 0. Beginning on line 53, we loop over the four control intervals; each pass through this loop integrates a single control interval over which there is a constant inducer level. On line 57 we loop over the `sample_pattern` array, such that the loop variable `num_stps` will contain the number of integration steps that need to be incremented to get to the next observation time point. Because there are three candidate observation time points within each control interval, this loop will repeat three times for each control interval; note that the time between observations has been chosen to be non-uniform, as set by the `sample_pattern` array. In line 57, the time stepping iterates over the specified number of steps using `step_func`. In the time stepping loop, we apply the RK4 step function in line 59 to the current state vector in `current_state` with the prescribed inducer level specified by `inducer_vector[interval]`. The step counter is incremented in line 60, to track the number of RK4 steps taken and thus the time elapsed. In line 62 we store the state vector, which at this point corresponds to an observation time point; the time is computed and recorded in line 63.

This manual implementation of the simulation is complex, but it provides significant control over the numerics and the OED problem structure. For example linear or parameterized spline-based inputs can be defined over each control interval and the candidate sampling schedule can be customized to accommodate any experimental constraints. Note that the OED algorithm will select which time points should be used for observation: here the user is pre-specifying candidate times to be considered.

```python
63 #merge all inducer levels into a single inputs vector
64 inputs = cs.vertcat(initial_inducer,inducer_vector)
65 # create list for observation structure
66 observation_list = []
67 #create list to store response names
68 observ_names, observ_type, observ_times = [], [], []
69 # loop over samples (time points)
70 for i in range(len(sample_list)):
71   #create a unique name for mrna and prot samples
72   mrna_name = 'mrna_'+'t'+"{0:0=2d}".format(times[i])
73   prot_name = 'prot_'+'t'+"{0:0=2d}".format(times[i])
74   #create mean and var tuple for mrna and prot observ.
75   mrna_stats = cs.vertcat(sample_list[i][0], 0.005)
76   prot_stats = cs.vertcat(sample_list[i][1], 0.005)
77   #create casadi function for mrna and prot stats
78   mrna_func = cs.Function(mrna_name,[inputs,parameters],[mrna_stats])
79   prot_func = cs.Function(prot_name,[inputs,parameters],[prot_stats])
80   #append the casadi function and distribution type to obs struct
81   observation_list.extend([(mrna_func,'Normal'), (prot_func,'Normal')])
82   #store observation names, useful for plotting
83   observ_names.extend([mrna_name,prot_name])
84   #store observation type
85   observ_type.extend(['RNA','Prot'])
86   #store observation time
87   observ_times.extend([times[i]]*2)
```

Listing 9: Code for assembly of the `observ_list` argument for use in the `Model` class constructor call for the dynamic gene expression model.

The result of the code in Listing 8 is a list, `sample_list`, containing CasADi symbols for each state variable at the candidate observation time points. The NLoed model constructor accepts CasADi functions for each observation variable and therefore these functions must now be created for each state and time point. The type of probability distribution for each state variable observation also needs to be specified. In our example we assume a normal distribution with constant variance. In Listing 9 a loop is used to construct CasADi functions for each state and time point, each of which is then stored in an observation list for passage to the NLoed `Model` constructor. In line 65 we concatenate the initial inducer level, `initial_inducer`, and the four control interval inducer levels in the `inducer_vector` to form the overall set of four inputs for the time series. In line 67, the `observation_list` list is declared and in lines 69 several lists for the observation name, species type and time point are initialized; these are needed to provide NLoed with observation names and for organizing data during analysis. On line 71, the loop for building up the observation list begins; this loop iterates over each observation state symbol in the sample list. In lines 73-74, each state's observation is given a unique string name, marking its species type and time. On lines 76-77, the observation's statistics – mean and variance – are concatenated together. In this example we assume all observations have a fixed variance of 0.005. On lines 79-81 we construct CasADi functions mapping the four experimental inputs (inducer levels) and the model parameters to the observation statistics. On line 82, we append each CasADi function in tuples with the `Normal` label, to the observation list. (The `Normal` label tells CasADi that each observation is assumed to be normally distributed and to expect functions for the mean and variance.) In lines 86 and 88 we store the auxiliary information about the species type and time point for downstream plotting and data.

```
88 #list the inpit and parameter names
89 input_names =
   ↪ ['Init_Inducer','Inducer_1','Inducer_2','Inducer_3','Inducer_4']
90 parameter_names =
   ↪ ['log_Alpha','log_K','log_Delta','log_Beta','log_Gamma']
91 #instantiate the model object
92 model_object = nl.Model(observation_list, input_names,
   ↪ parameter_names)
```

Listing 10: Code for the creation of an NLOED `Model` object for the mRNA-protein dynamic model.

Finally, after generating the observation structure in the previous loop, we are ready to create the NLOED model. Shown in Listing 10, in lines 90 and 91, we declare the input and parameter names and on line 93 we create an NLOED model object named `model_object` using the NLOED `Model` class constructor. This model object can be used in the same way as the one created for the CcaS/CcaR system in the main text; for a more complete description of features see the NLoed's Github repository at `https://github.com/ingallslab/NLoed`.

### 5 Timing Comparison with `aceBayes`

To provide readers with a general idea of the computational trade-offs between a local tool like NLoed and Bayesian methods such as those implemented in `aceBayes`, we performed a simple timing analysis for the model presented in the main text. All timing analysis was performed on a 2019 Macbook Air with a 1.6 GHz Intel Core i5 processor and 8GB of memory. We used Python's `time` package and the function call `time()` to record the time taken to generate the optimal design, via `nl.Design()` in Listing 4 of the main text. This was recorded as taking 0.245 seconds.

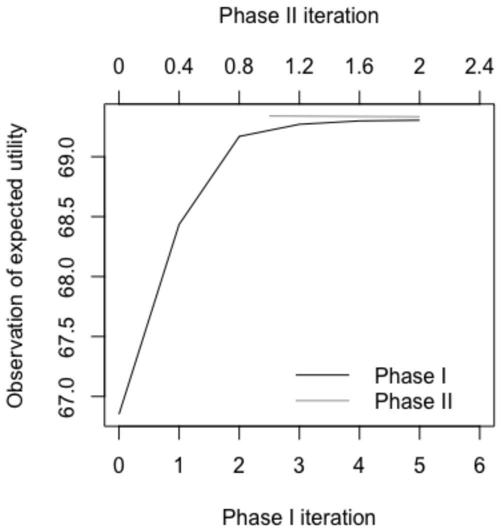


Figure 3: Convergence diagnostic plot generated for the `aceBayes` design optimization.

We then implemented the same model in `aceBayes`, as shown in Listing 11. The optimal design generated by NLoed is generated conditionally, based on both the past parameter estimates from the initial dataset, `data0`, as well as the previous design information, `design0`. The `aceBayes` package does not allow for this type of iterative conditional design. Instead, we used an analogous procedure by providing a prior mean and variance of the parameter estimates computed from `data0` and using this to generate the optimal Bayesian design. To compute the prior mean and variance for the initial data, `data0`, we used the parameter estimates as mean and the covariance matrix generated via NLoed's `fit()` and `evaluate()` call in Listing 3 of the main text. The numerical values of these estimates are given in supplementary Listing 11. We chose to compare NLoed to `aceBayes` using a Bayesian D-optimal objective as this was the most similar method to NLoed's locally D-optimal objective. We chose to use Monte Carlo as the computational method (see the `aceBayes` documentation for a complete listing of options). As `aceBayes` uses a stochastic algorithm to perform the design optimization, convergence must be verified. To perform a fair comparison we selected the minimum number of iterations so that the visual diagnostic generated by a call to `plot` for the `acenlm` object indicated no further increase in the objective value. The convergence diagnostic plot is shown in Figure 3. We found this to be 5 iterations of `aceBayes`'s first phase. (The `aceBayes` package has a two-phased optimization algorithm; the first for coarse but rapid improvement of the objective, and the second for fine-tuning.) We additionally performed two iterations of the second phase, the minimum required to confirm that there was no increasing trend in the objective provided by the fine-tuning in the second phase. To record the time taken to generate the Bayesian optimal design, we used `aceBayes`'s internal timing functionality by printing the `time` field of the `acenlm` object. We found the Bayesian optimal design with 5 iterations of the first phase, and two iterations of the second phase took 19.179 seconds.

```
library(acebayes)
library(MASS)
#Set seed for reproducible results
set.seed(1)
#Set sample size, as in main text
n<-15
#Define a Latin Hyper Cube starting design
start.d<-matrix(24 * randomLHS(n = n, k = 1), nrow = n, ncol = 1,
                dimnames = list(as.character(1:n), c("u")))
#Define the prior's mean and variance to match that computed from
#design0 using NLoed's fitting algorithm
#NOTE: This is done to replicate NLoed's conditional design
mu = c(6.31506051, 9.15833531, 0.88530341, 2.14426291)
sigma = matrix(c(
    0.00112157, -0.00021089,  0.00120845, -0.0004012,
        -0.00021089,  0.00083638, -0.0007035, 0.0008058,
        0.00120845, -0.0007035, 0.0041213, -0.00207542,
        -0.0004012, 0.0008058,  -0.00207542,  0.00189346),4,4)
#Define a function for the prior
prior_mc<-function(B){
  out<-mvrnorm(n = B, mu, sigma)
  colnames(out)<-c("ln_A0", "ln_A", "ln_N","ln_K")
  out}
#Create the D-optimal Bayesian design using Monte Carlo
D_MC<-acenlm(formula = ~ (exp(ln_A0) +
    exp(ln_A)*u^exp(ln_N)/(exp(ln_K)^exp(ln_N) + u^exp(ln_N))),
             start.d = start.d, prior = prior_mc,
             N1 = 4, N2 = 2, B = c(1000, 1000),
             lower = 0, upper = 100, method = "MC")
#Plot the convergence information to ensure convergence
plot(D_MC)
#Print the time
D_MC$time
#Print the optimal design
D_MC$phase2.d
```

Listing 11: Code for implementing the main text model and a Bayesian D-optimal design using the `aceBayes` package in `R`.

Our tesing indicates that the time cost of optimization for `aceBayes` is approximately two orders of magnitude slower than NLoed (.245 seconds vs 19.179 seconds). For small models such as that presented in the main text, the computational burden of either approach is minimal, and so `aceBayes` may be preferred – especially given the additional utility that the Bayesian analysis can provide in hedging against uncertainty. However if the user wishes to explore a variety of model formulations and design constraints, and then assess their performance via simulation study, the additional computational cost of a Bayesian approach may be limiting. The same trend may bear out when considering more complex models, as the computational cost scales with the number of inputs and parameters. We conclude that NLoed and `aceBayes` are complementary tools. For experiments that may involve rapid iteration between design and measurement, or situations where modelers wish to explore many model and design variants in large simulation studies, NLoed is likely to provide better scalability. This may be common in systems biology studies involving cell or bacterial cultures where costs are lower and iteration is more feasible. Bayesian approaches, in contrast, are ideal for more expensive one-shot experiments, perhaps involving human or animal subjects, where it is important to hedge against uncertainty as much as possible in the design phase.
